# The impact on authors and editors of introducing Data Availability Statements at Nature journals

**DOI:** 10.1101/264929

**Authors:** Rebecca Grant, Iain Hrynaszkiewicz

**Author notes:** Correspondence should be addressed to Rebecca Grant, Springer Nature, The Campus, Trematon Walk, Wharfdale Road, London N1 9FN.

## Abstract

This paper describes the adoption of a standard policy for the inclusion of data availability statements in all research articles published at the Nature family of journals, and the subsequent research which assessed the impacts that these policies had on authors, editors, and the availability of datasets. The key findings of this research project include the determination of average and median times required to add a data availability statement to an article; and a correlation between the way researchers make their data available, and the time required to add a data availability statement. This paper will be presented at the International Digital Curation Conference 2018, and has been submitted to the International Journal of Digital curation.

## Introduction

In September 2016, it was announced that all research papers accepted for publication in Nature, and the life science journals in the Nature family, would be required to include information on whether the data underpinning their study were made available, and how others can access it. This followed a successful 2-month pilot implementation of the policy at five Nature journals between March 2016 and May 2016: Nature Neuroscience, Nature Physics, Nature Communications, Nature Cell Biology and Nature Geoscience. The introduction of data availability statements (DAS) by Nature journals aligned Nature's policies with a standardised Springer Nature research data policy framework, which was introduced earlier in 2016 (Hrynaszkiewicz et al, 2017).

DASs are written by authors to provide information on where the data supporting the results reported in their article can be found, and if and how they can be obtained. Although there is no mandated format for these statements, template examples are provided to Nature authors, including:

- The datasets generated during and/or analysed during the current study are available in the [NAME] repository, [PERSISTENT WEB LINK TO DATASETS].
- The datasets generated during and/or analysed during the current study are available from the corresponding author on reasonable request.
- All data generated or analysed during this study are included in this published article (and its supplementary information files).

## Analysing Data Availability Statements at Nature Journals

Two phases of research were undertaken to analyse the impact of this new policy. The aim of this research was to assess the ways by which researchers chose to make their data available, and to measure the additional time required by editors and production staff to add data availability statements to manuscripts (which gives an indication of cost to the publisher).

For the five pilot journals, editors were asked to self-report the number of additional minutes it took to process a manuscript to ensure an appropriate DAS was provided. For Nature Communications - one of the five pilot journals, and which publishes a large number of articles - editors self-reported an overall average time for all papers they were responsible for over the pilot period, rather than reporting the time required for every manuscript. As the data were collected with different methodology and with lower precision they are not included in this analysis. For all other journals, editors recorded a time (in minutes) to add a DAS for every paper that they were responsible for in the pilot period. Copy editors and production staff were also asked to provide an estimated average additional manuscript processing time for all papers they handled in this initial period. Editors, production and copy editing staff were also invited to provide comments on the process of providing the DAS.

Once the papers were accepted for publication, the text of each DAS was read and categorised into one of four different types according to a coding mechanism created for the project, by Iain Hrynaszkiewicz (Head of Data Publishing, Springer Nature). This was done for four journals (Nature Neuroscience, Nature Physics, Nature Cell Biology and Nature Geoscience), including 82 papers, by assigning the author’s description of their data availability to one of the four categories/codes. The code that best matched the main message of the DAS or was applicable to the majority of data referred to in the DAS was used. The codes used were the following:

- Type 1 stated that the data is available from the author on request.
- Type 2 stated that the data had been included in the manuscript or its supplementary material.
- Type 3 stated that some or all of the data is publicly available, for example in a repository.
- Type 4 stated that figure source data was included with the manuscript. This is a method of data sharing used by some authors in a subset of Nature journals that publish life sciences research. Some journals encourage authors to provide the source data behind their figures/plots as spreadsheets. It is specific to the Nature journals and is not mandated and, as such, is relatively uncommon, but was important to capture in this analysis for internal purposes.

Univariate and bivariate analysis was then used to interpret the coded data. For the same four journals, it was found that it took 10 minutes extra editorial time on average, or a median time of 8 minutes per paper, to add a DAS (Figure 1). Additionally, 5 minutes extra copy editing time was required. For Nature Communications, 90% of editors reported 15 minutes or less to add a DAS on average. Overall, the addition of the DAS had an impact of approximately 15-20 minutes editorial and production time per accepted paper across all five journals in the 2-month pilot.

When the average time to add a DAS is presented by type of code statement, rather than by journal, the type 1 statement, where data are available on request, was the fastest, on average (5.9 minutes) to add to a paper. The type 3 statement, where some or all data are publicly available, took the most amount of additional time (18.2 minutes), in the four pilot journals included in this analysis.

**Figure 1.**
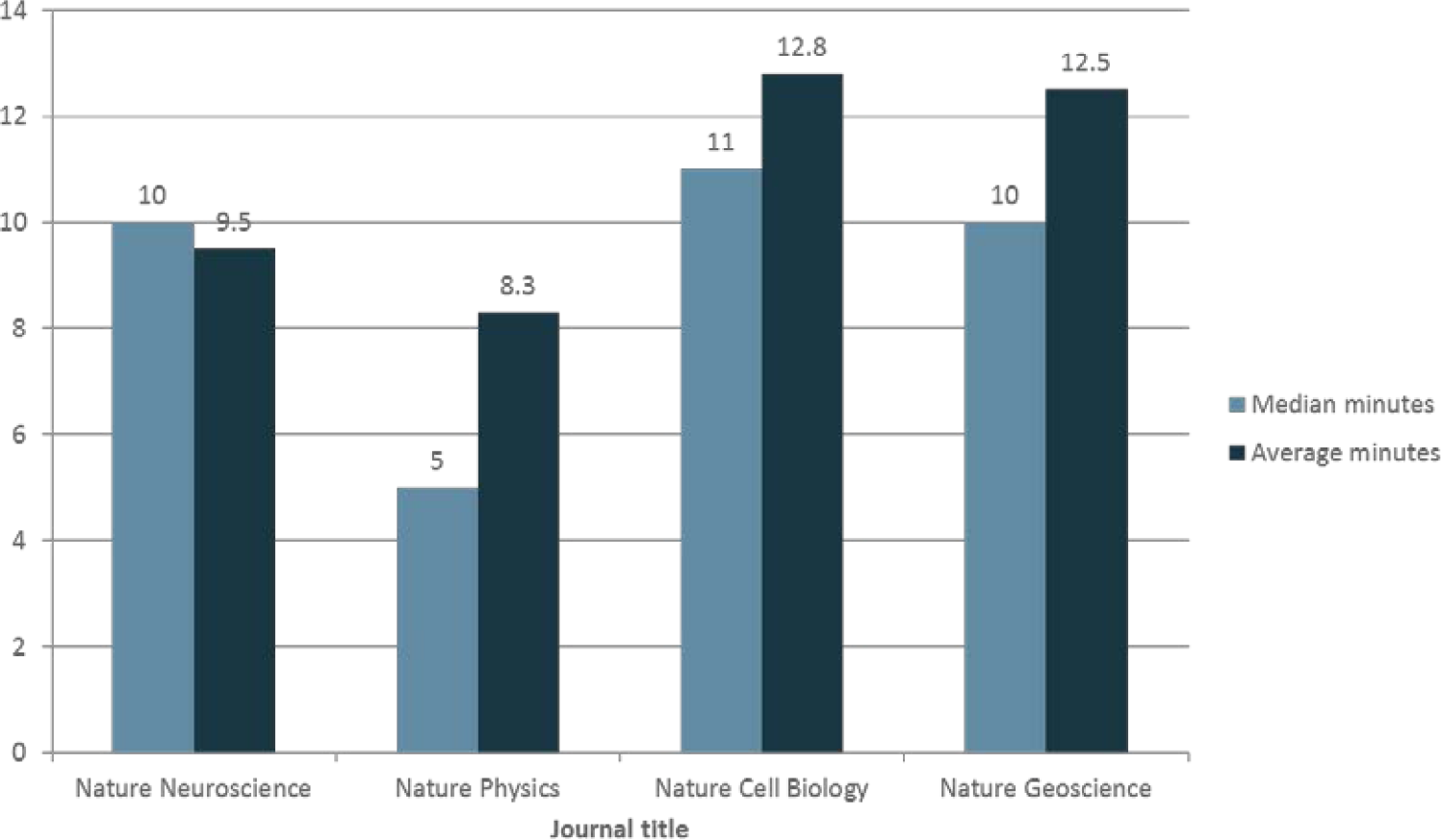
Median and average editor time to add DAS by journal (minutes).

**Figure 2.**
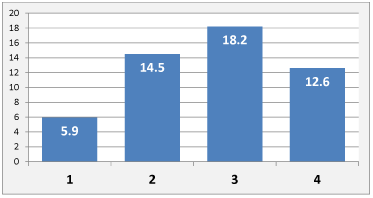
Average time by statement type in minutes (n=82).

The second phase of this project gathered data using the same methodology from an additional 20 journals. These were from the biological and physical sciences, which introduced the same DAS policy as the previous journals, from September 2016. These journals provided the same information as the previous five, including the DAS, and time required to add the DAS. Data were gathered by each journal for 2 months after implementation of the policy, between September 2016 and February 2017. Because the time data relied on editor self-reporting, there are a proportion of manuscripts for which time data were missing (n=112). However, coding of the text of the data availability statements, by Hrynaszkiewicz and Rebecca Grant (Research Data Manager, Springer Nature), for all accepted manuscripts in the study period was carried out.

In this larger sample of papers, average editor time to add a DAS decreased for all types of statement compared to the pilot journals (which was 10 minutes on average). In this second analysis (Figure 3), which pooled all data from the initial pilot journals and the additional 20 journals, the type 1 statement remained the fastest (3.4 minutes on average) to add to a paper and the type 3 statement remained slower (average 6.2 minutes) than type 1 and type 2 (3.7 minutes average). The type 4 statement took the longest to implement in the second analysis. It decreased only marginally (average 12.5 minutes) for the type 4 statement but there were the fewest (10) of these statements and papers recorded.

**Figure 3.**
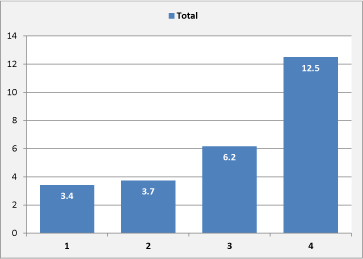
Average time by statement type in minutes (n=329).

As several journals included in the analysis published low numbers of papers requiring a DAS in the 2-month period after implementation of the policy, and to provide a more useful analysis of statement types for 25 journals, statement type data are reported by the journals’ major subject discipline rather than by journal (Figure 4).

**Figure 4.**
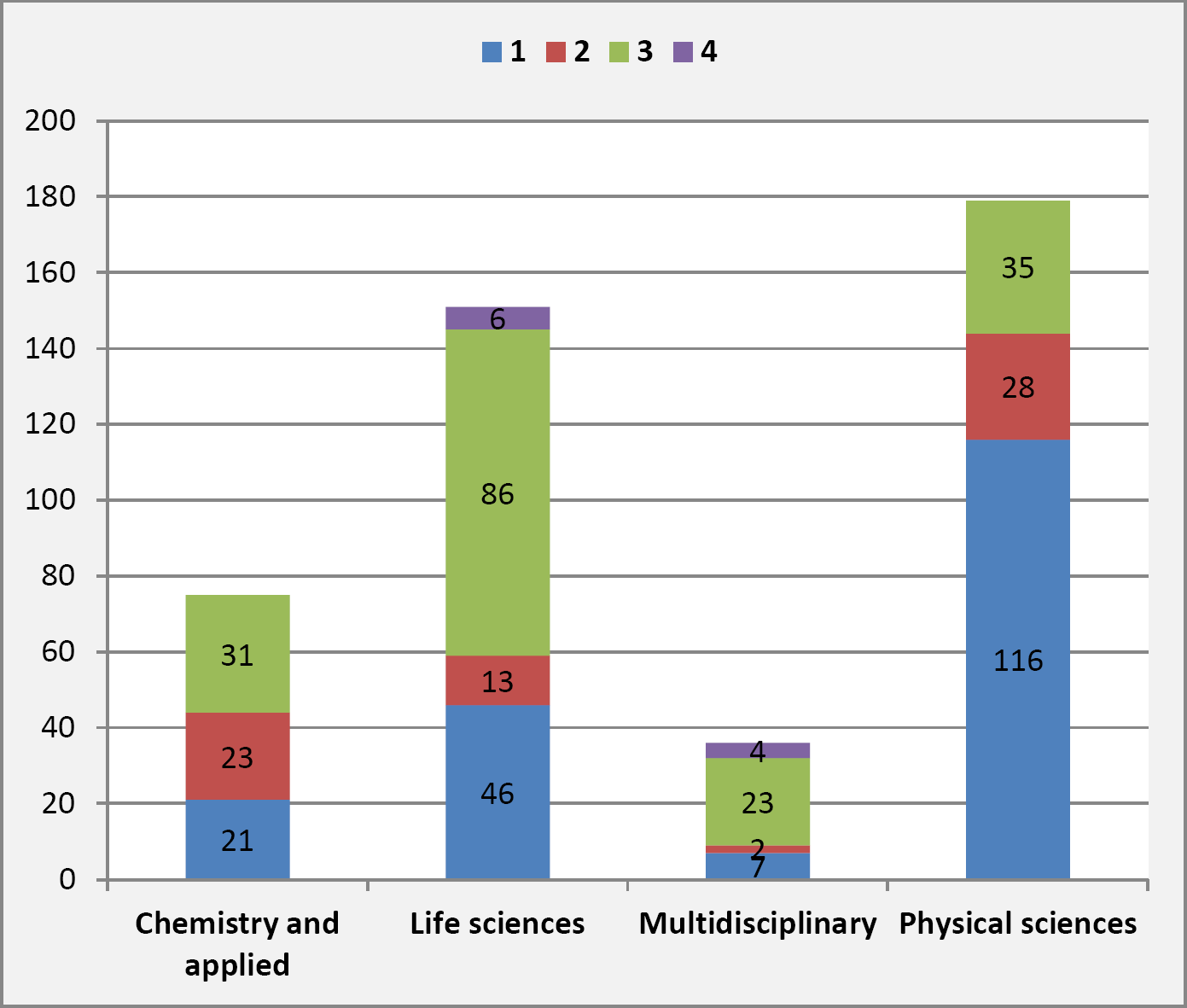
Distribution of statement types by journal’s major discipline (n=441).

The classification of journals and papers into four broad subject groups provides some indication of different data sharing practices in different disciplines, For example, in life sciences there was the highest proportion of papers that made supporting data available in a repository (86/151 papers; 57%). In physical sciences the proportion of papers making data available in a repository was the lowest (35/179 papers; 20%).

## Limitations

Data relating to time was self-reported by around 50 editors and support staff and so may not always be reported consistently, and is subject to individual biases and interpretations of their time. Where no time-related data were provided, those papers were removed from the analysis. Where an editor reported zero minutes or “negligible” these were recorded as 1 minute. The coding method was developed by Hrynaszkiewicz and coding carried out by Hrynaszkiewicz and Grant. Where there was disagreement in coding, this was resolved by consensus.

## Discussion and Conclusion

Adding mandatory DASs to all accepted articles in journals operated by professional editors increases manuscript processing time. The publisher deemed the time added to be reasonable in the context of total manuscript submission to publication time given the importance and benefits of including a DAS. Indeed, the additional time did not limit implementation of mandatory DASs at all Nature journals in 2016, after implementation at the five pilot journals.

The type 3 DAS, where data are made publicly available, took longer to introduce to papers than statements where data were available on request or declared as being available with the supplementary information files. The type 3 statements include the most non-standard text, as well as links to be verified, which likely caused the longer processing time. This type of statement - making data available with, and linked to published articles - is anticipated to provide the most benefit to authors and readers, such as the potential for greater numbers of citations to those papers (Dorch at al, 2015, Piwowar et al, 2013, Sears, 2014).

The second, larger, group of journals to introduce mandatory DASs reported that fewer additional minutes were added to the editorial time needed to process a manuscript. Possible reasons for this include greater editor and author awareness of the policy and supporting documents; improved internal communication and editor training after the pilot; greater attention being needed on the pilot journals, which informed, and made more rapid, editor training on handling future DAS for manuscripts in their discipline.

The findings of the study demonstrate the impact of data availability statements on editorial staff and the journal publication workflow, and the need to consider increased manuscript processing requirements when mandatory DAS are introduced. As well as understanding the benefits of increasing accessibility to research data is also important to understand costs - particularly for publishers, funding agencies and policy makers. Information from this analysis has informed the selection of data policies by other Springer Nature journals. It has also informed the development of in-house administrative support for academic editors, so that journals without professional editors can also introduce DASs consistently. Simple, practical information, such as additional minutes to process manuscripts, is valuable for editors and support staff in understanding the impacts of editorial policy changes.

The disciplinary differences in data sharing practices indicated in Figure 4 are likely due to larger numbers of community repositories being available in life sciences, combined with long standing data sharing mandates for life sciences communities, which are enforced in the editorial process at the Nature journals. In physical sciences, there are fewer mandates and the analysis also includes Nature Physics, where, in high energy physics for example, data produced and analysed can be very large - produced at large central facilities - making sharing of data online challenging. In such cases “data available on reasonable request” can be a pragmatic choice.

Although there is an increase in the time required by editorial staff to process articles, the introduction of DAS will also have repercussions for other data curation stakeholders. The incorporation of standardised DASs is likely to become more widespread across journal publishers. Since Springer Nature began introducing standardised data policies, similar initiatives have been introduced by other large publishers such as Elsevier and Wiley. The standardisation of research data policies for journal publishing is being progressed by initiatives such as the Research Data Alliance's Data Policy Standardisation and Implementation Interest Group. There are also discipline-specific initiatives to standardise and harmonise journal and publisher research data policy, in chemistry and high energy physics and medicine (Taichman et al, 2017). There is growing attention on the provision of mandatory DASs from publishers, institutions and funding agencies, particularly in the UK where DASs are a requirement of the Research Councils UK (RCUK) Common Principles on Data Policy. DASs are simple and interoperable - between stakeholders, publishing platforms and research disciplines - mechanism for communicating the availability of supporting data which can aid monitoring of compliance with data policies of funders, publishers and research communities. They can also support funder policies which require data publication by providing evidence of compliance through descriptions of publicly accessible datasets.

The increase of prevalence of DASs by journal publishers necessitates support and training for researchers and editors to enable researchers to share and cite their data wherever possible.

Increased prevalence of DASs will also enable further research, using machine-driven approaches (such as with natural language processing, text and data mining), across multiple journals and publishers, to analyse the types of DASs provided - and types of data sharing practiced - by researchers in different disciplines and journals. Further research would also be welcome on associations, if any, between the provision of particular types of data availability statement and research visibility and impact as studies have tended to be limited to specific disciplines and journals (Rowhani-Farid and Barnett, 2016).

## Data availability

A partially anonymised dataset that supports the figures and graphs in this paper is available in figshare with the identifier: https://doi.org/10.6084/m9.figshare.5809617.

In this shared dataset some data columns have been removed, such as personal comments made by authors and editors during correspondence about inserting a DAS, as we do not have consent to publish them. Internal manuscript identifiers have been replaced with ascending numbers. Requests for additional data or for support with reusing the data should be emailed to the authors, or to

## Acknowledgements

The authors thank Christos Petrou and Graham Smith at Springer Nature for support with data analysis; Sowmya Swaminathan, Head of Editorial Policy at Nature Research for input into policy implementation at the Nature journals; and all editors and administrative staff at the Nature journals who contributed to data acquisition.

